# SARS-CoV-2 convergent evolution cannot be reliably inferred from phylogenetic analyses

**DOI:** 10.1101/2021.05.15.444301

**Authors:** Yoon-Seo Jo, Asif U. Tamuri, Greg J. Towers, Richard A. Goldstein

## Abstract

A homoplasy is a trait shared between individuals that did not arise in a common ancestor, but rather is the result of convergent evolution. SARS-CoV-2 homoplasic mutations are important to characterise, because the evidence for a mutation conferring a fitness advantage is strengthened if this mutation has evolved independently and repeatedly in separate viral lineages. Yet detecting homoplasy is difficult due to insufficient variation between sequences to construct reliable phylogenetic trees. Here, we develop a method to more robustly identify confident homoplasies. We derive a maximum likelihood (ML) tree, with taxa bearing seemingly recurrent mutations dispersed across the tree, and then, for each potentially homoplasic mutation, we derive an alternative tree where the same taxa are constrained to one clade such that the mutation is no longer homoplasic. We then compare how well the two trees fit the sequence data. Applying this method to SARS-CoV-2 yields only a few instances where the constrained trees have significantly less statistical support than unconstrained tree, suggesting phylogenetics can provide limited support for homoplasy in SARS-CoV-2 and that caution is needed when inferring evidence of convergent evolution from phylogenetic methods in the absence of evidence from other sources.

## Introduction

Control of the COVID-19 pandemic has been hampered by the emergence of multiple SARS-CoV-2 variants of concern (VOC). The British B.1.1.7 (Rambaut *et al*., 2020), South African B.1.351 (Tegally *et al*., 2020), and Brazilian P.1 (Faria *et al*., 2021) VOC lineages arose in late 2020 and quickly became dominant within the countries in which they were first detected. The Indian B.1.617.2 variant has also recently been designated as a VOC (Public Health England, 2021). Additional variants under investigation (VUI), which may be later classified as VOC, continue to appear. There is evidence that some of these new strains have evolved resistance to immunity acquired by prior infection or current vaccines (Greaney *et al*., 2021; Thomson *et al*., 2021; Wang *et al*., 2021) possibly undermining development of vaccine induced herd immunity (Naveca *et al*., 2021). VOC mutations likely also increase transmissibility (Davies *et al*., 2021) and virulence or severity of disease. Hence, the surveillance of emerging variants, and the identification of those most likely to have altered phenotypes, is paramount in understanding and controlling the pandemic.

Sequence changes naturally arise due to stochastic errors in replication. Most of these are deleterious and are rapidly eliminated from the virus population. Some changes may represent an adaptation or response to selective pressure, and might increase in frequency due to the resulting increase in fitness. Current identification of VOCs is based principally on these increases in infection frequency. However, rapid increases in mutation frequency can occur in the absence of any adaptive change due to founder effects (Díez-Fuertes *et al*., 2021), poor public health or increased sampling in the region in which the variant emerged, circulation in a sub-population more prone to infection or ‘hitchhiking’ onto an advantageous change elsewhere (Smith and Haigh, 1974). Ambiguity in the reason for increases in frequency highlights the importance of other indications of a fitness advantage. Furthermore, we want to determine which variants represent threats to public health prior to significant virus spread.

Confidence in the significance of an observed mutation increases if we observe the same mutation independently occurring in separate SARS-CoV-2 lineages. Viruses of disparate lineages that experience the same selective pressure in a common niche (e.g., host humoral immunity from vaccination or replication in a new host), will potentially alight on similar solutions (e.g., mutations that change the epitope, or adapt to human host interactions). These independently recurring changes are known as homoplasies, or examples of convergent evolution. For these reasons, homoplasic changes in the SARS-CoV-2 evolution are important to reliably identify.

Unfortunately, identification of homoplasies is not trivial. Identification of homoplasic changes generally relies on a reliable phylogenetic tree which establishes that lineages with the same mutation are actually independent. This is difficult for SARS-CoV-2 due to the limited genetic diversity (Duchene *et al*., 2020) and therefore low phylogenetic signal. Also, phylogenetic trees are mathematical models representing a simplified reconstruction of complex real-life events, with parameters optimised to best fit the data at hand. These models are too simple to represent the underlying evolutionary process and tree construction depends on robustness of the phylogeny to the inevitable model misspecification. Typically, a large number of phylogenetic trees approximately equally fit the observed sequence data. Lack of phylogenetic signal greatly increases the size of the plausible tree set, whilst model misspecification alters the trees included in this set and their relative likelihood values. In this case, the tree with the highest likelihood (the ‘best-fitting’ tree) is selected, more or less at random, from an immense set of plausible trees with nearly equal support (Morel *et al*., 2020), which makes it extremely unlikely that this “best” tree represents the true evolutionary history. It is difficult to conclude that some feature of the evolutionary dynamics, such as homoplasy, is supported by the phylogeny if the set of plausible trees includes trees that do and do not support that feature.

Of course, failure to detect homoplasy via phylogenetics does not necessarily mean that a particular change is not homoplasic; rather it means the statistical support for homoplasy, as it is observed on a phylogenetic tree, is weak. It is also important to recognise that lack of homoplasy does not necessarily indicate that the change in sequence is not adaptive or does not have important phenotypic consequences. Similarly, homoplasies will arise with some frequency even in the absence of adaptive change. Finally, we note that each position is considered independently when estimating the support for homoplasy. This is important because phenotypes, such as vaccine resistance, may require multiple changes.

Phylogenetics is just one source of information. The speed of phylogenetic inferences compared to, for example *in vitro* or *in vivo* laboratory experiments make evolutionary analyses extremely useful and attractive for selecting the sequence changes for further study using these more time-consuming methods. To gain robust phylogenetic insights and prioritise SARS-CoV-2 mutations for phenotypic or experimental analyses, we must have a measure of confidence in phylogenetic output. Specifically, we need to address how confident an inference derived from the best-fitting tree is, given the large selection of plausible alternatives. One possibility is to examine whether there are plausible trees where there is no homoplasy at the position under study.

Here, we evaluate the statistical confidence for inferring homoplasy from phylogenetic trees. We first construct the best maximum likelihood (ML) tree derived from SARS-CoV-2 sequences, from which potential homoplasies are identified. Then, for each observed potential homoplasy, we derive an alternative tree, which is the best possible (maximum likelihood) phylogeny given a specific topological constraint. Theoretical examples are provided in Supplementary Fig. 1a and 1b. The topological constraint in each case is a tree in which all taxa bearing the potentially homoplasic mutation are manipulated to be monophyletic. In this latter tree, the mutation is no longer homoplasic, as only one change from a common ancestor would give rise to daughter taxa all with the same change. We then ask: is the constrained tree a significantly worse fit for the sequence data? To determine this, we use the Approximately Unbiased (AU) test (Shimodaira, 2002), which ascertains whether a certain phylogenetic structure is supported by the sequence data. If the best constrained tree is indeed significantly less well supported, then this analysis provides evidence for homoplasy. In contrast, if the data are also found to be consistent with the constrained tree topology wherein the homoplasy has been removed, then the given site has weak phylogenetic support for homoplasy.

## Results

Using a dataset of 3947 SARS-CoV-2 sequences (downloaded 5^th^ February 2021 from https://www.gisaid.org/), we first identified potential homoplasies on the unconstrained ML tree. These were defined as instances where there were multiple, independent changes to the same amino acid (AA) (Methods). This returned 1697 potential homoplasies, to be further examined. We then made a series of maximum likelihood trees constrained to eliminate each homoplasy in turn. We assumed a null hypothesis where the best constrained tree did not provide a significantly worse fit to the sequence data than the unconstrained ML tree. We conducted the AU test and calculated the false discovery rate (FDR), which controls for the fraction of false predictions of homoplasy.

Strikingly, this analysis produced only three strongly supported (FDR < 0.05) homoplasies, Δ69/70 and Δ143 in the Spike protein and P309S in nsp2 (Fig. 1; Table 1). One mutation, H78Y in ORF3a, had moderate support (0.05 ≤ FDR < 0.10), and 15 sites had weak support (0.10 ≤ FDR < 0.20) for homoplasy (Fig. 1). The remaining 1678 sites failed to reject the null hypothesis under 0.20 FDR. That is, the constrained trees fit to the sequence data were not significantly worse than the unconstrained ML tree.

**Figure 1.**
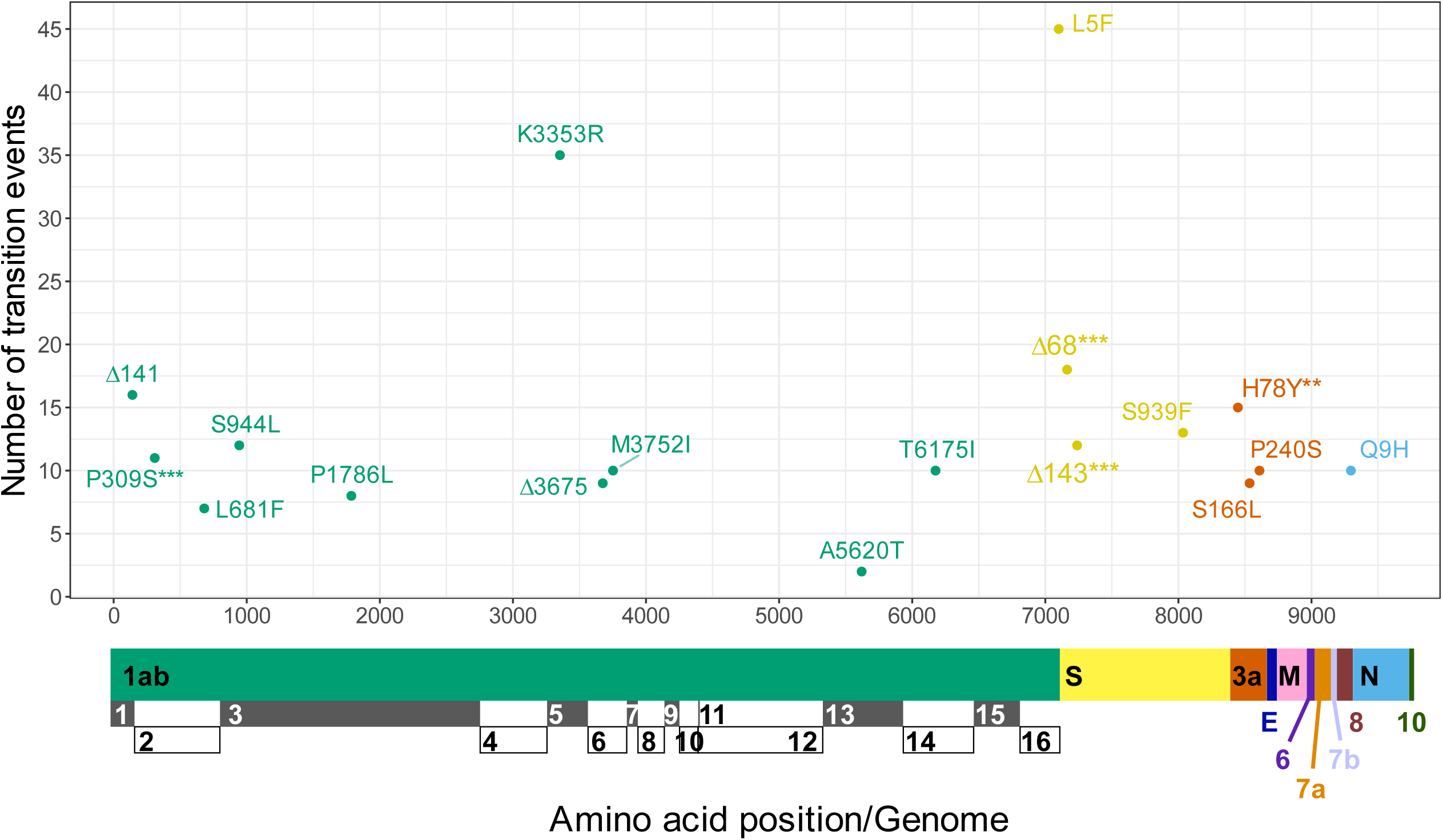
Supported homoplasies. Coding mutations with *** strong (FDR < 0.05), ** moderate (0.05 ≤ FDR < 0.10), or weak (0.10 ≤ FDR < 0.20) support for homoplasy are located on the SARS-CoV-2 genome, along with the total number of each mutational event occurring in the alignment. The non-coding deletion at nucleotide position 28271 is not shown. Black and white boxes denote nsp in ORF1ab polyprotein.

**Table 1.** Mutations with support for homoplasy. FDR column: *** strong support (FDR < 0.05); ** moderate support (0.05 ≤ FDR < 0.10); * weak support (0.10 ≤ FDR < 0.20) for homoplasy. The equivalent positions of mutations in polyprotein ORF1ab are: P309S = nsp2 P129S; L681F = nsp2 L501F; S944L = nsp3 S126L; P1786L = nsp3 P968L; K3353R = nsp5 K90R; Δ3675 = nsp6 Δ106; M3752I = nsp6 M183I; A5620T = nsp13 A296T; T6175I = nsp14 T250I. “nc28271del” is a deletion in a non-coding region, at nucleotide position 28271 of the genome.

| Gene | Mutation | *p*-values<br>(uncorrected) | FDR |
| --- | --- | --- | --- |
| Spike | $\Delta 68$ | 6.70E-07 | ***0.003 |
| Spike | $\Delta 143$ | 2.25E-06 | ***0.006 |
| ORF1ab (nsp2) | P309S | 2.16E-05 | ***0.036 |
| ORF3a | H78Y | 4.84E-05 | **0.060 |
| N/A | nc28271del | 1.37E-04 | *0.135 |
| ORF1ab (nsp6) | M3752I | 1.87E-04 | *0.154 |
| Spike | S939F | 2.21E-04 | *0.156 |
| ORF1ab (nsp1) | $\Delta 141$ | 2.97E-04 | *0.163 |
| ORF1ab (nsp6) | $\Delta 3675$ | 2.71E-04 | *0.163 |
| ORF1ab (nsp3) | S944L | 3.39E-04 | *0.166 |
| ORF3a | S166L | 3.70E-04 | *0.166 |
| Nucleocapsid | Q9H | 4.97E-04 | *0.197 |
| ORF1ab (nsp14) | T6175I | 6.62E-04 | *0.197 |
| ORF1ab (nsp3) | P1786L | 5.27E-04 | *0.197 |
| ORF1ab (nsp5) | K3353R | 5.69E-04 | *0.197 |
| ORF3a | P240S | 6.30E-04 | *0.197 |
| Spike | L5F | 6.79E-04 | *0.197 |
| ORF1ab (nsp13) | A5620T | 7.64E-04 | *0.199 |
| ORF1ab (nsp2) | L681F | 7.52E-04 | *0.199 |

Of the 19 homoplasies with support (FDR < 0.20), 5 were deletions and 14 were nonsynonymous base changes. Of all the changes in the protein sequences in our dataset (n=4940), 161 were insertions/deletions (indels) and 4779 were base changes. Thus, a greater proportion of indels (3.1%) than base changes (0.29%) were supported for homoplasy in our analysis. However, we propose that this is due to an innate bias in the tree search process which systematically gives indels greater confidence for homoplasy than nucleotide changes, as discussed below.

## Discussion

We have assessed the use of phylogenetics to detect homoplasic mutations in SARS-CoV-2, and demonstrated that reliable inference of homoplasy is challenging. Specifically, we evaluated support for homoplasic mutations by considering whether an unconstrained ML tree provided a significantly better fit to the sequence data than the best constrained tree where each homoplasy was eliminated. Of the 1697 mutations that appeared homoplasic on the unconstrained tree, only 19 had an FDR less than 0.20.

We identified a number of mutations with statistical support for homoplasy (Fig. 1; Table 1). Of these changes, only four were observed in the Spike protein. There is a paucity of information regarding the functional consequences of the non-Spike mutations highlighted by our approach, such as P309S (equivalently P129S) in nsp2, M3752I (equivalently M183I) in nsp6, H78Y in ORF3a, and Q9H in Nucleocapsid. Δ141-143 in the ORF1ab nsp1 C-terminus has been proposed to hinder nsp1 gene expression inhibition and interferon antagonism (Benedetti *et al*., 2020). However, these preliminary analyses require additional molecular and structural studies, and warrant further investigations of the SARS-CoV-2 accessory proteins.

By contrast, Spike mutations have been scrutinised because Spike is an important target for the adaptive immune response, the viral antigen used in all current vaccines, and is relatively simple to functionally characterise *in vitro*. The statistically supported homoplasies Spike Δ69/70 and Δ144, are lineage-defining deletions in VOC B.1.1.7 (Rambaut *et al*., 2020). Δ69/70 is associated with higher viral loads and may explain the increased transmissibility of B.1.1.7 (Kemp *et al*., 2021), while Δ144 abolishes binding of the neutralising antibody, 4A8, and thus is expected to alter Spike antigenicity in the population (McCarthy *et al*., 2021). None of the other Spike defining mutations found in VOC B.1.1.7, B.1.351 or P.1 (O’Toole *et al*., 2020; Rambaut *et al*., 2020; Tegally *et al*., 2020; Faria *et al*., 2021) were significantly supported for homoplasy by our analysis (Supplementary Table 1). This was surprising, because recurrent VOC mutations are widely believed to be homoplasic (Ferrareze *et al*., 2021; Lemmermann *et al*., 2021; Martin *et al*., 2021; Resende *et al*., 2021). For instance, the Spike receptor binding domain (RBD) mutations E484K, which reduces neutralisation by sera (Collier *et al*., 2021) (Fig. 2), and N501Y, which increases affinity to ACE2 (Starr *et al*., 2020) (Fig. 3), had weak support for homoplasy. Despite the acquisition and appearance of common VOC mutations in different lineages (Fig. 2a; Fig. 3a), we showed that we can still generate a plausible phylogenetic tree that suppresses the homoplasy (Fig. 2b; Fig. 3b). In other words, while N501Y, E484K, and other key recurrent VOC mutations Spike L452R, Spike S477N, Spike H655Y, Spike Q677H and Spike P681H (Supplementary Figs. 2-6) may be homoplasic by other lines of evidence, the lineages bearing these mutations can reasonably form a single clade, and thus phylogenetic tree construction does not provide support for their convergent evolution.

**Figure 2.**
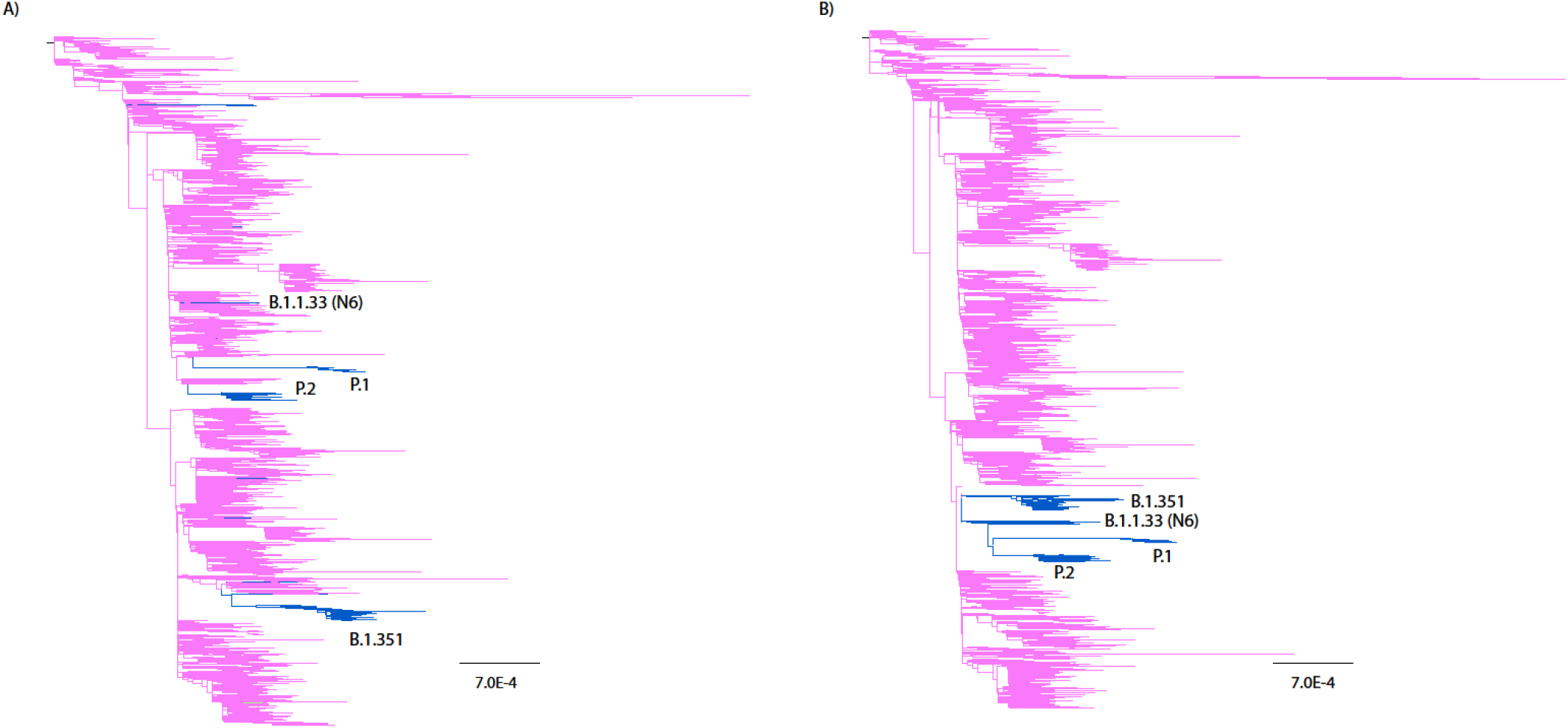
Phylogenetic trees colour coded to depict amino acids at position 484 of Spike (magenta: Glutamic acid, blue: Lysine). (A) Unconstrained ML tree; (B) Alternative tree optimised under the constraint that the lysines are monophyletic, and the E484K mutation is not homoplasic. Alternative tree was not significantly worse at matching sequence data (FDR = 0.338).

**Figure 3.**
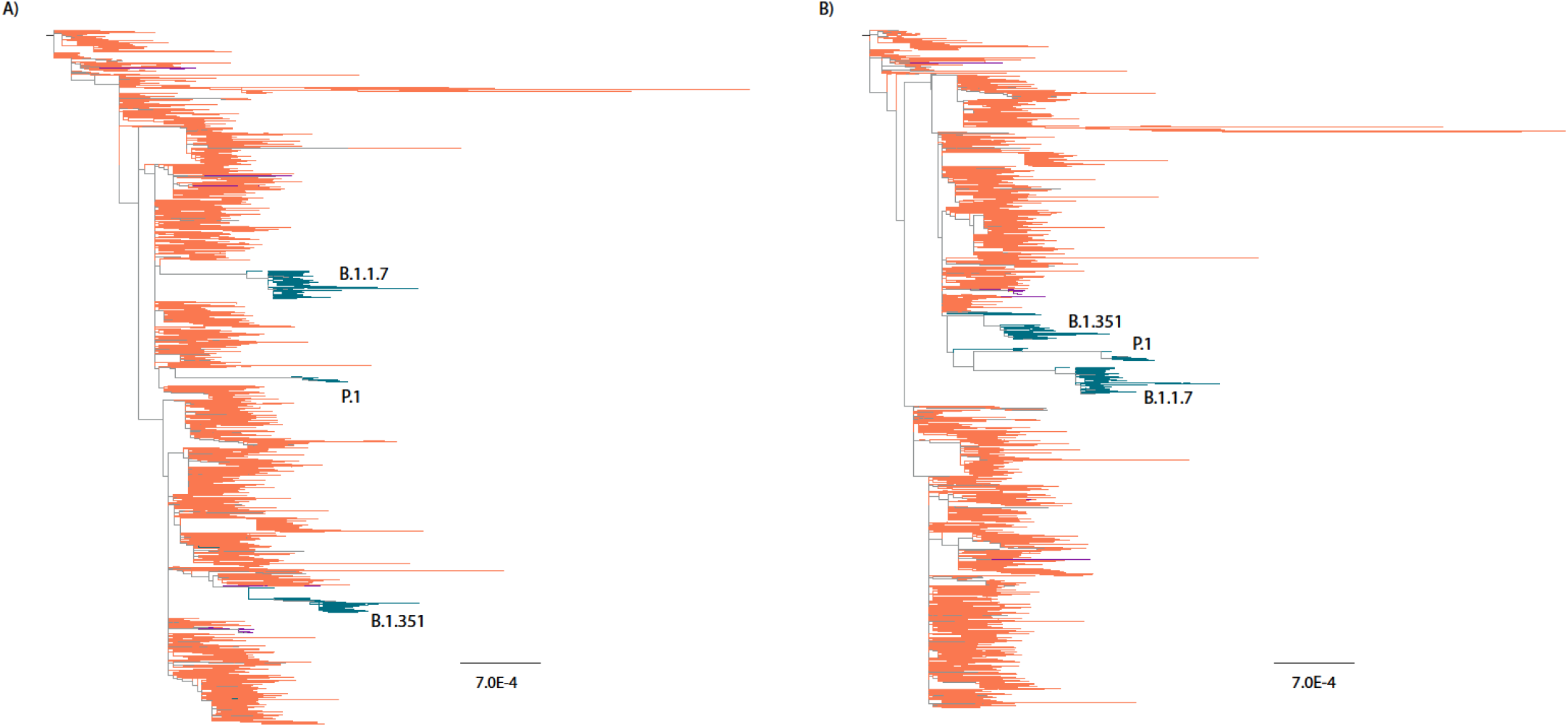
Phylogenetic trees colour coded to depict amino acids at position 501 of Spike (orange: Asparagine, green: Tyrosine). (A) Unconstrained ML tree; (B) Alternative tree optimised under the constraint that the tyrosines are monophyletic, and the N501Y mutation is not homoplasic. Alternative tree was not significantly worse at matching sequence data (FDR = 0.618).

We did observe statistical support for two other homoplasic mutations in the Spike protein: S939F and L5F (Fig. 1). L5F has been preferentially observed in specific sequencing laboratories, suggesting that it might be a sequencing artefact (De Maio *et al*., 2020; Turakhia *et al*., 2020). Likewise, S939F was reported to be a recurrent artefact by De Maio *et al*. (2020), although other studies have predicted that this mutation, which occurs in the ‘fusion core’ of the HR1 domain, may destabilise the Spike protein (Ahamad *et al*., 2021), including the post-fusion assembly (Cavallo and Oliva, 2020).

The lack of statistical support for homoplasy for the vast majority of mutations results from the general lack of phylogenetic signal in SARS-CoV-2 sequences, as has been noted previously (Morel *et al*., 2020). In addition, several other factors may play an important role.

SARS-CoV-2 phylogenies generally contain long branches (Figs. 2, 3), consistent with an expanding viral population. The presence of these long branches complicates phylogenetic analyses. Firstly, the places where these long branches connect to the rest of the tree are poorly resolved and can be moved around liberally without a significant statistical forfeit (Mossel and Steel, 2005). Secondly, even with maximum likelihood analyses, there will be a systematic bias in tree reconstructions due to long branch attraction, in which clades with long branch lengths are inferred to be closely related, regardless of their true evolutionary relationship (Felsenstein, 1978). Finally, it is clear that the assumptions of the phylogenetic analysis are not fulfilled. For instance, the assumption that every site in the alignment can be treated independently is contradicted by observations of epistatic interactions (Kemp *et al*., 2021). Errors due to model misspecification are likely to be a greater problem with longer branches, as the pattern of mutations will be less constrained. As a result, in rapidly expanding viral populations, in which longer branches are more common, there will be an intrinsically high level of phylogenetic uncertainty (Turakhia *et al*., 2020).

Importantly, such problems may be particularly important for SARS CoV-2 analyses, as many of the VOC variants have arisen on long branches (Fig. 2, 3). This may result from a number of different factors. Long branches may reflect evolution in an unsampled reservoir, or accelerated sequence change due to relaxed selective constraints, increased mutation rates due to changes in the polymerase or RNA structures, or adaptive bursts in atypical (e.g., immunodeficient) hosts. While the specific reasons for these long branches are of obvious public health relevance, they increase the phylogenetic uncertainty, including the identification of the specific mutations leading to new VOCs.

We noted above that the fraction of indels that are homoplasic is significantly higher than the fraction of base changes. It is not immediately obvious why Spike Δ69/70 and Δ144, which are found on a long-branch leading to VOC B.1.1.7, were not subject to the same uncertainties as other long-branch changes. The answer might lie in how phylogenetic analyses treat indels and nucleotide changes differently. ML calculations aim to estimate the best parameter values (e.g., topology, branch lengths) to maximise the probability of the sequence data given the model (tree). Homoplasies involve multiple changes of the same type, which is less likely than a single change, decreasing the likelihood of a phylogenetic tree. Algorithms ameliorate this reduction by resolving/removing the homoplasy, but this often has the adverse effect of creating new homoplasies – in other words, by resolving one homoplasy, we often produce a homoplasy at a different site. Thus, maximal likelihood resolves homoplasies which have a substantial penalty in favour of homoplasies that incur a lesser penalty. The problem with indels is that they are considered to be ‘missing data’, such that they do not have any likelihood consequences. Therefore, phylogenetic algorithms preferentially resolve homoplasies in base substitutions at the detriment of increasing homoplasies in indels. This may produce a general bias towards identifying homoplasic indels with high but potentially mistaken confidence.

It is important to reiterate that failure to demonstrate confident homoplasy via phylogenetics does not mean a mutation is not homoplasic or causing a significant phenotypic change. Conversely, even if a mutation is homoplasic, that does not mean that it has phenotypic consequences. Non-adaptive convergence is much more frequent in closely related lineages than would be predicted by standard substitution models, especially so in sites which have recently diverged (Castoe *et al*., 2009; Goldstein *et al*., 2015), as in the case of the circulating SARS-CoV-2 lineages.

The survey of all potential homoplasic mutations required substantial computational resources. As a result, we analysed only a subset of the available sequences, those selected by Nextstrain (https://nextstrain.org/sars-cov-2). Statistical support for homoplasy was substantially larger when the analysis was repeated with a smaller data set of 524 taxa. The decrease in support with increasing sequence data can result from a number of factors. Fewer sequences results in longer branches and fewer constraints on substitution model parameters, causing biases in the tree reconstruction due to long branch attraction and model misspecification. These biases can easily result in the appearance of seemingly well supported but erroneous homoplasies. In addition, the inclusion of additional taxa exacerbates the difficulties and uncertainties inherent in tree reconstruction, making the set of plausible phylogenetic trees substantially larger and more likely to include trees consistent with absence of homoplasy. For these reasons, it is unlikely that an analysis of a larger number of taxa would yield greater support for the homoplasic mutations. The time required for these calculations precluded the analysis of the most recent data. In particular, our analysis was performed prior to the observation of the various Indian B. 1.617 variants. We would expect that the overall conclusion, that phylogenetics provides little support for potential homoplasies, would not be affected. Determining the homoplasies for which there is support would require continued updating of the analysis, but can be streamlined by neglecting homoplasies without statistical support and where additional sequences do not result in additional mutations in the updated unconstrained tree.

Many studies employ phylogenetic methods to derive conclusions of convergent evolution, which are error-prone for the reasons described above. Furthermore, it is not uncommon to locate the occurrence of mutations in different lineages based on an estimated phylogenetic tree, without further assessment of the confidence for the tree itself (van Dorp, Richard, *et al*., 2020; van Dorp, Tan, *et al*., 2020; Hodcroft *et al*., 2021). Our results demonstrate that inference of SARS-CoV-2 homoplasy from such a phylogenetic analysis is unreliable. Hence, we propose that to reliably infer homoplasy from a phylogenetic tree, it must be validated with a suitable statistical analysis. In contrast, we identify a mutation as homoplasic when the ensemble of plausible trees does not contain trees where that mutation is not homoplasic. As a result, we are still able to robustly evaluate statistical support for homoplasies in the absence of a well-supported phylogenetic tree, since our approach does not depend on knowing the ‘correct’ phylogeny.

## Methods

### Data acquisition

We confined ourselves to a subset of 3947 sequences assembled by Nextstrain in their global sample (https://nextstrain.org/sars-cov-2/, downloaded 5 February 2021). Corresponding full-length human SARS-CoV-2 sequences were retrieved from the Global Initiative on Sharing All Influenza Database (GISAID) (https://www.gisaid.org/), which they align to the reference Wuhan sequence using MAFFT. The alignment was further adjusted by hand using AliView v1.26 (Larsson, 2014).

### Phylogenetic inference

A maximum-likelihood (ML) phylogeny was constructed using IQ-TREE (v2.1.2) (Minh *et al*., 2020) under the substitution model proposed by Hasegawa *et al*. (1985) with empirical state frequency and gamma distribution of among-site rate variation (HKY+F+G). The tree was rooted using the Wuhan reference sequence (GISAID EPI_ISL_402125) as an outgroup. This was repeated ten times, and the ML tree with the highest log likelihood (LL) was chosen for subsequent analyses. We assigned sequence changes to branches on the phylogenetic tree by identifying differences in the consensus reconstructed ancestral states between adjacent nodes. Changes in nucleotide sequences were then translated to changes in protein sequences.

### Homoplasy identification and resolution

Homoplasies were identified by multiple changes to the same amino acid along different branches at a given site in a protein sequence, independently of the prior amino acid. For each identified homoplasy, we constructed a constraint tree in the form ((*a*_1_,*a*_2_,*a*_3_…),(*b*_1_,*b*_1_,*b*_3_…)), where (*a*_1_,*a*_2_,*a*_3_…) included the taxa downstream of any of the homoplasic changes that had the resulting amino acid; and (*b*_1_,*b*_1_*,b*_3_…) comprised taxa that were not downstream of any of these changes and had a different amino acid. Sequences with other than standard amino acids at that site or that were downstream of a homoplasic change but did not have the resulting amino acid were not contained in the constraint tree. Sequences that were downstream of multiple transition nodes were ignored. These constraints corresponded to resolving the homoplasy by confining the sequence change to the single branch leading to the taxa containing the corresponding sequence change. We then used IQ+TREE (HKY+F+G) to find the maximum likelihood tree given the constraints.

### Statistical analysis

The AU test (Shimodaira, 2002) as implemented in IQ-TREE was used to determine whether the constrained, alternative tree topology was a significantly worse fit to the data than the unconstrained ML tree topology, for each homoplasic site. False discovery rates (FDRs) were calculated using the Benjamini & Hochberg (1995) procedure of multiple hypothesis testing, including all observed amino acid changes and indels (n=4940). P values for mutations not observed to be homoplasic on the unconstrained tree were set to 1. Tests with FDR < 0.05 were categorised as having ‘Strong’, 0.05 ≤ FDR < 0.10 as ‘Moderate’, and 0.10 ≤ FDR < 0.20 as ‘Weak’ support for homoplasy. Trees were viewed on ChromaClade (Monit *et al*., 2019) and FigTree v1.4.4 (http://tree.bio.ed.ac.uk/software/figtree/).

### Analysis of indels

Treatments of indels are notoriously difficult, as they violate the assumption that each site can be analysed independently. To overcome these obstacles, we made a catalogue of all observed insertions and deletions relative to the reference strain, to include indels in non-coding regions. We then created a separate multiple sequence alignment encoding the presence or absence of each unique indel, encoded as binary data. When the site of a given insertion or deletion was covered by a subsequent deletion, the presence of the initial indel was represented as missing information. We used a simple Jukes-Cantor (JC) model (Jukes and Cantor, 1969) to re-create the ancestral states of the indels, and analysed the statistical support for homoplasy in the manner described above.

## Acknowledgements

GJT is funded by Wellcome Investigator Award 220863 and the G2P-UK National Virology consortium (MRC/UKRI grant MR/W005611/1). RAG is funded by UK Biotechnology and Biological Sciences Research Council Grant BB/P007562/1. We are grateful to Tom Peacock (Imperial College London) and Florencia Tettamanti Boshier (University College London) for helpful discussions.

