## Supplementary figures for "SARS-CoV-2 convergent evolution cannot be reliably inferred from phylogenetic analyses"

(a)

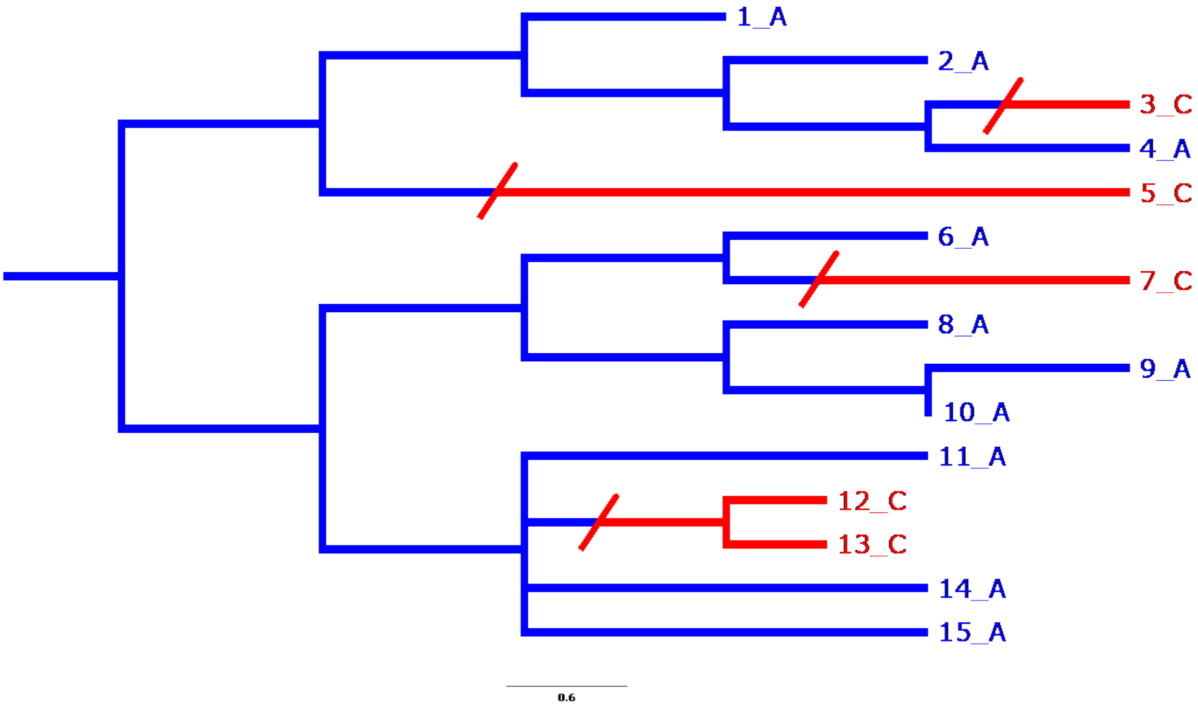


(b)

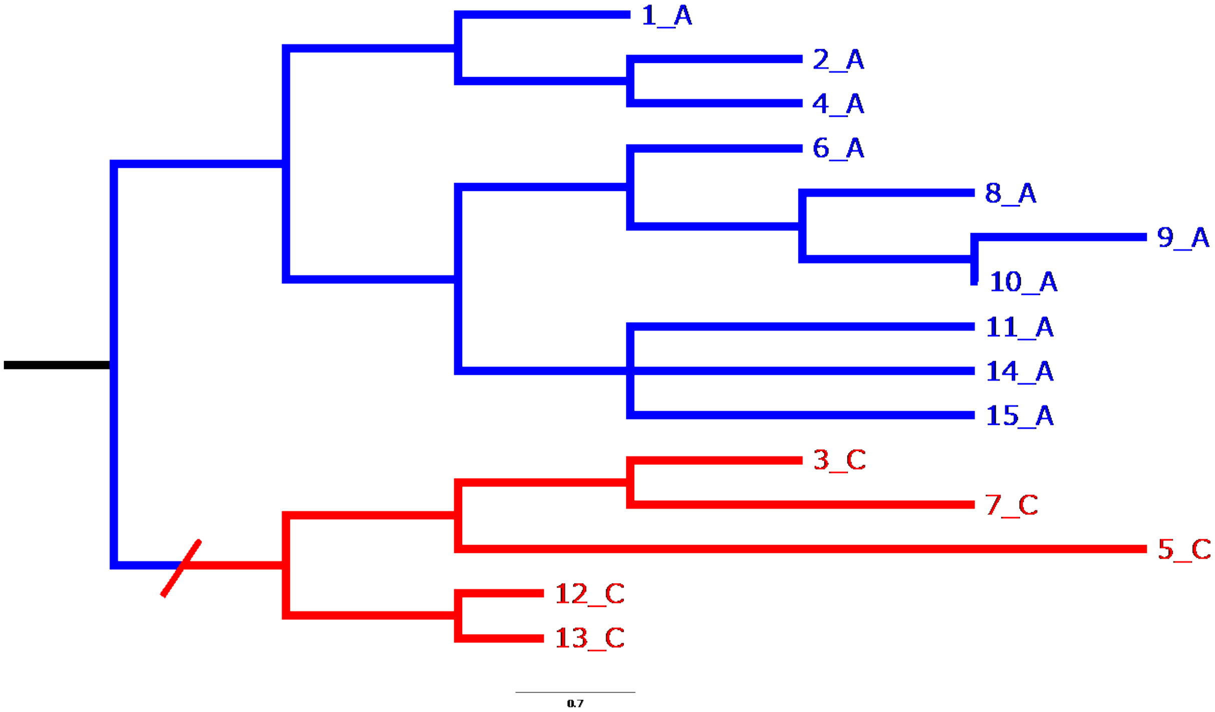


**Supplementary Figure 1. Exemplar trees.**

Taxa are labelled in the form *'x_y'* where *x* is the taxa name, and *y* is the amino acid at a particular site. (a) An unconstrained tree, akin to a tree inferred by maximum likelihood (ML) without topological manipulation. Here, there are four distinct changes from amino acid A to amino acid C. (b) A constrained alternative tree. All taxa with the 'homoplasic mutation' (amino acid C) are brought together to one clade, thus forming a constraint. The Newick tree specified in the reconstruction of this alternative tree would be: ((1_A,2_A,4_A,6_A,8_A,9_A,10_A,11_A,14_A,15_A) , (3_C,5_C,7_C,12_C,13_C)). Each test involves comparing the unconstrained ML tree (a), and the best tree inferred with the constraint (b), to determine if the constrained tree has significantly less statistical support than the unconstrained tree.

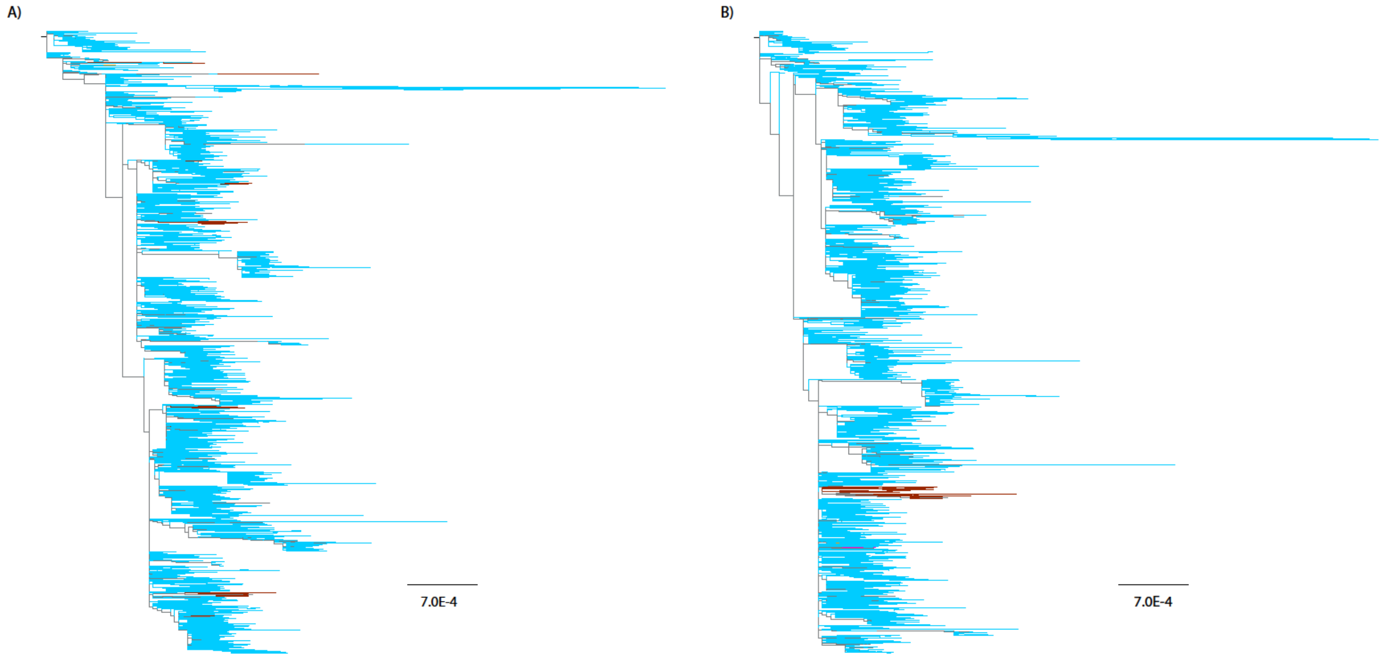


**Supplementary Figure 2. Spike L452R**
Trees depict amino acids at position 452 of Spike, with taxa coloured by the amino acid present at this site (blue: Leucine, maroon: Arginine). Sequences are applied to (a) the unconstrained ML tree topology and (b) the alternative tree topology wherein the homoplasic taxa with Arginine are constrained together (FDR = 0.301). Wuhan-1 (EPI_ISL_402125) was used to root all trees.

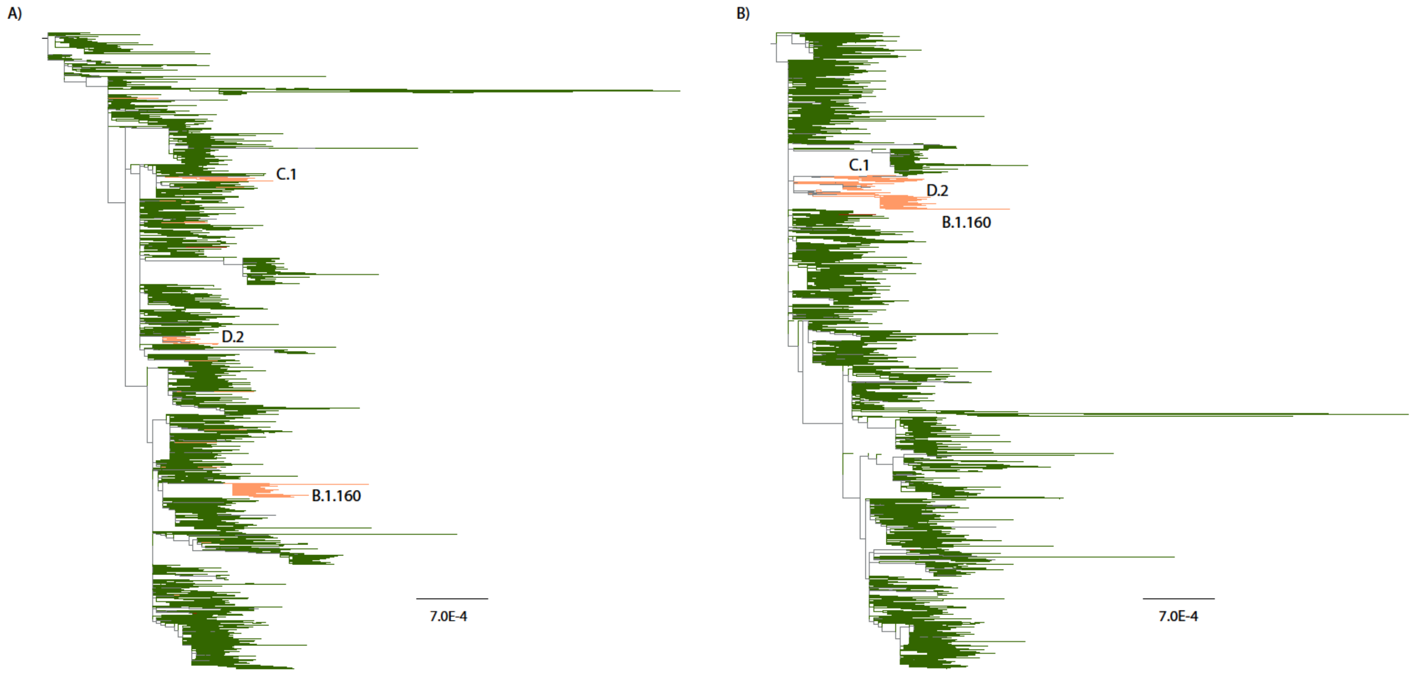


**Supplementary Figure 3. Spike S477N**
Trees depict amino acids at position 477 of Spike, with taxa coloured by the amino acid present at this site (green: Serine, orange: Asparagine). Sequences are applied to (a) the unconstrained ML tree topology and (b) the alternative tree topology wherein the homoplasic taxa with Asparagine are constrained together (FDR = 0.344). Wuhan-1 (EPI_ISL_402125) was used to root all trees.

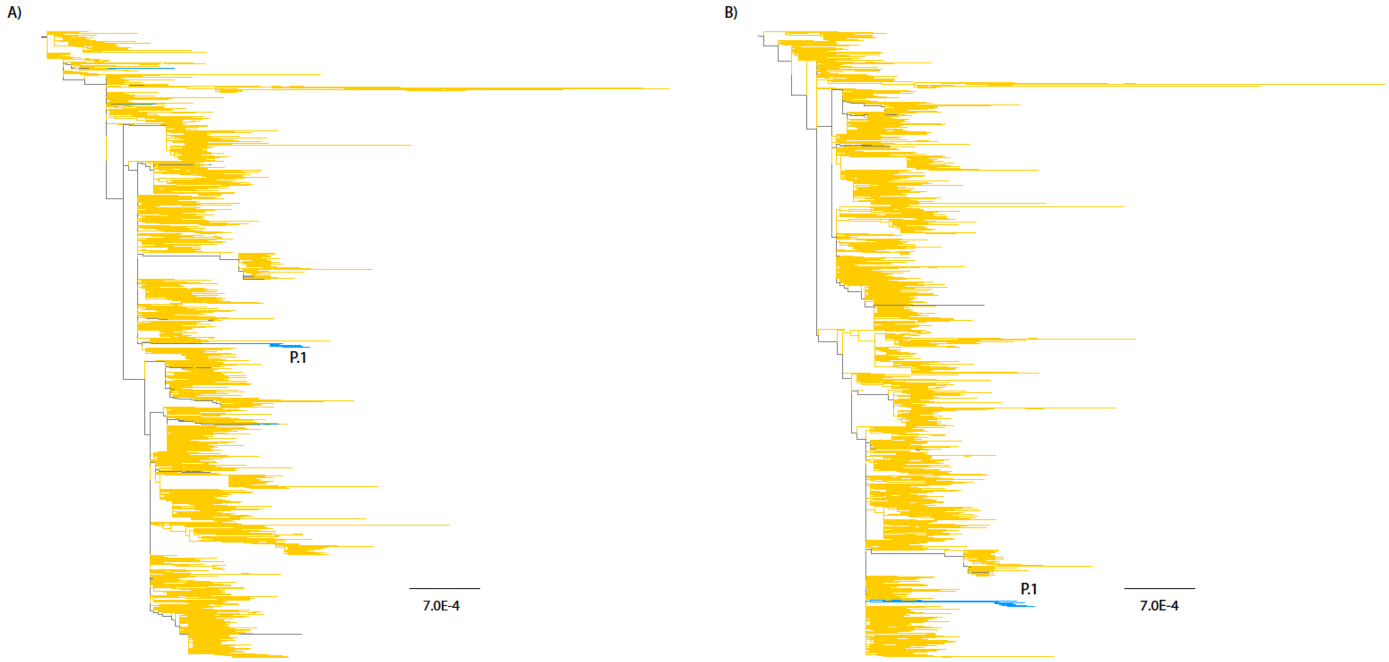


**Supplementary Figure 4. Spike H655Y**

Trees depict amino acids at position 655 of Spike, with taxa coloured by the amino acid present at this site (gold: Histidine, blue: Tyrosine). Sequences are applied to (a) the unconstrained ML tree topology and (b) the alternative tree topology wherein the homoplasic taxa with Tyrosine are constrained together (FDR = 0.270). Wuhan-1 (EPI_ISL_402125) was used to root all trees.

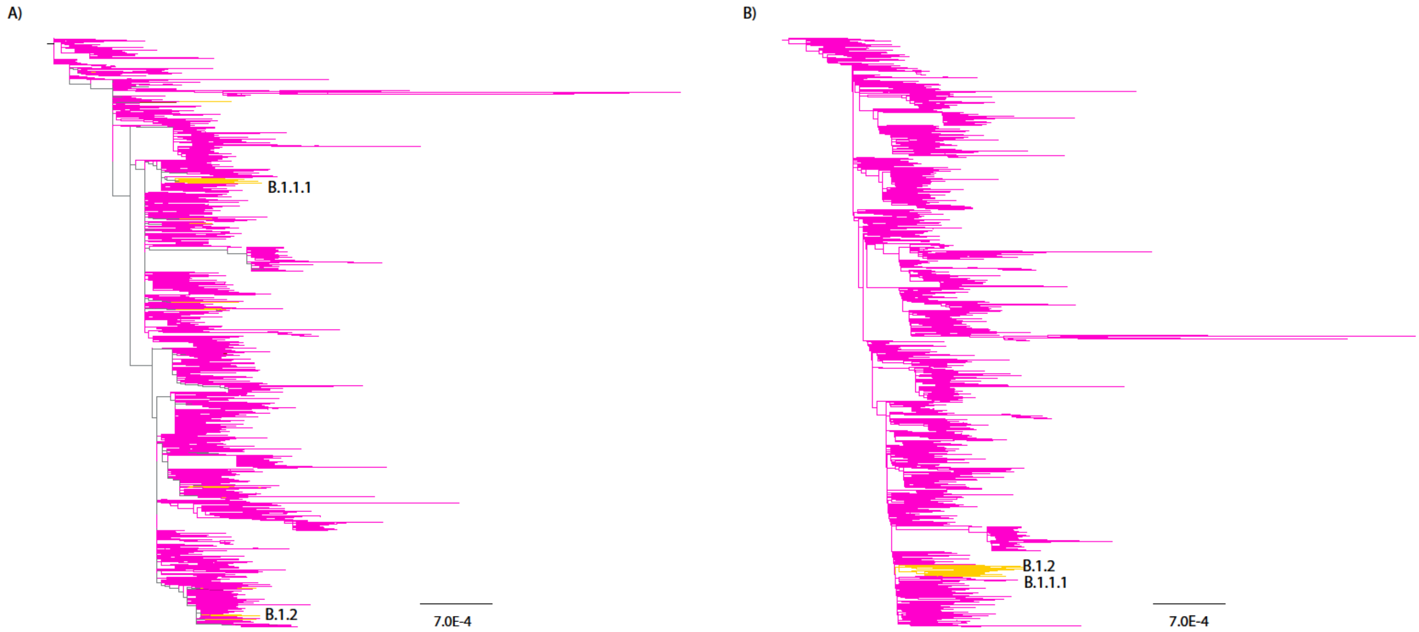


**Supplementary Figure 5. Spike Q677H**

Trees depict amino acids at position 677 of Spike, with taxa coloured by the amino acid present at this site (magenta: Glutamine, yellow: Histidine). Sequences are applied to (a) the unconstrained ML tree topology and (b) the alternative tree topology wherein the homoplasic taxa with Histidine are constrained together (FDR = 0.267). Wuhan-1 (EPI_ISL_402125) was used to root all trees.

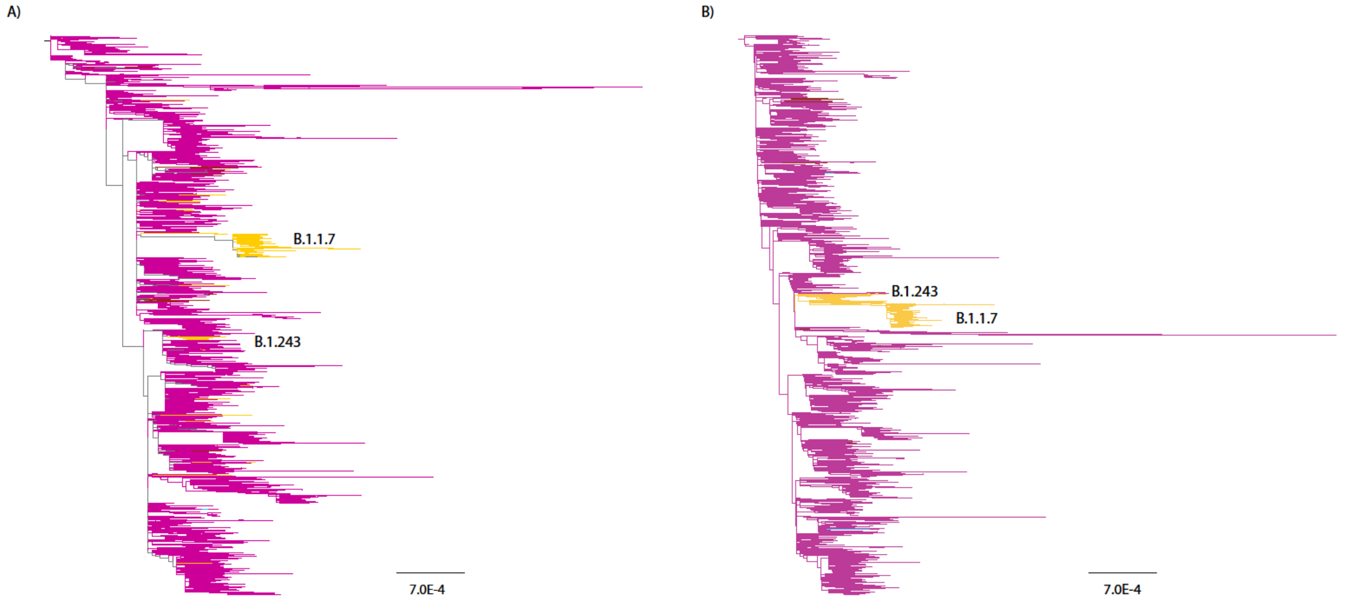


**Supplementary Figure 6. Spike P681H**
Trees depict amino acids at position 681 of Spike, with taxa coloured by the amino acid present at this site (magenta: Proline, yellow: Histidine). Sequences are applied to (a) the unconstrained ML tree topology and (b) the alternative tree topology wherein the homoplasic taxa with Histidine are constrained together (FDR = 0.325). Wuhan-1 (EPI_ISL_402125) was used to root all trees.
